# AutoUnmix: an autoencoder-based spectral unmixing method for multi-color fluorescence microscopy imaging

**DOI:** 10.1101/2023.05.30.542836

**Authors:** Yuan Jiang, Hao Sha, Shuai Liu, Peiwu Qin, Yongbing Zhang

## Abstract

Multiplexed fluorescence microscopy imaging is widely used in biomedical applications. However, simultaneous imaging of multiple fluorophores can result in spectral leaks and overlapping, which greatly degrades image quality and subsequent analysis. Existing popular spectral unmixing methods are mainly based on computational intensive linear models and the performance is heavily dependent on the reference spectra, which may greatly preclude its further applications. In this paper, we propose a deep learning-based blindly spectral unmixing method, termed AutoUnmix, to imitate the physical spectral mixing process. A tranfer learning framework is further devised to allow our AutoUnmix adapting to a variety of imaging systems without retraining the network. Our proposed method has demonstrated real-time unmixing capabilities, surpassing existing methods by up to 100-fold in terms of unmixing speed. We further validate the reconstruction performance on both synthetic datasets and biological samples. The unmixing results of AutoUnmix achieve a highest SSIM of 0.99 in both three- and four-color imaging, with nearly up to 20% higher than other popular unmixing methods. Due to the desirable property of data independency and superior blind unmixing performance, we believe AutoUnmix is a powerful tool to study the interaction process of different organelles labeled by multiple fluorophores.

## 1 Introduction

Fluorescence microscopy imaging is an essential tool in biomedical research to obtain cellular tissue characteristics of samples. Labeling imaging samples with multiple fluorescent dyes may result in spectral overlap between emission signals detected by distinct channels, leading to cross-talk and blurring of signals, and thereby limiting the number of fluorophores that can be employed simultaneously^1^. To address this problem, various spectral unmixing algorithms have been developed for multispectral imaging to obtain the spectral features, and separate them into corresponding image sets of individual dyes, thus outputting pure sample images.

To identify the individual fluorescent dyes of biomarkers and the corresponding ratios of each pixel in each channel, researchers have proposed several spectral unmixing methods, which are mainly divided into linear unmixing and non-linear unmixing. Based on the assumption of linear mixing of emission fluorescence, various linear spectral unmixing methods usually require known emission spectral profiles of fluorophores. Linear unmixing^2^ can be formulated as the inverse problem of *Y* =*AX*, where *Y, X*, and *A* represent the mixed image, unmixed image, and mixing matrix respectively. Although linear unmixing is employed widely in multi-color microscopy imaging^3, 4^, the unmixing performance mainly relies on the quality of the calibrated emission spectra and the mixing matrices. When the calibrated environment is not consistent with the actual experimental environment, the measured emission spectra may be shifted significantly, affecting the unmixing results greatly^5^. Additionally, traditional iterative optimization methods to solve the inverse problem of linear unmixing suffer from heavy computational costs. To improve computational efficiency, some researchers have proposed unmixing methods using vertex component analysis (VCA)^6^ and independent component analysis (ICA)^7^. Meanwhile, based on the property that both the fluorescent intensity and the ratios of different channels are non-negative, the non-negative matrix factorization (NMF) method^8, 9^ has been proposed to solve the inverse problem.

In order to solve the problem of requiring prior spectral information and low computational efficiency, many non-linear spectral unmixing methods have been proposed in recent years. Seo et al.^10^ design an information theory-based iterative optimization method by minimizing mutual information between channels of mixed images to blindly unmix 15 colors without reference spectra. With the wide spread of machine learning, related methods have been applied in the field of fluorescent microscopy imaging. Some studies utilize support vector machine (SVM)^11^ to learn the pixel feature of fluorescent images and cluster them to realize spectral unmixing and separation of cellular locations^12, 13^. In addition, McRae et al.^14^ propose Learning Unsupervised Means of Spectra (LUMoS) without the need for prior measurements of their emission spectra and constraints on the number of fluorophores that can be utilized. This work uses K-Means to learn the spectral characteristics of individual fluorophores from mixed images to achieve blind spectral unmixing. However, the latter two machine learning methods require hand-craft iterating parameters and the settings of hyperparameters may impact the final effects of unmixing. Besides, methods based on phasor analysis^15^ show some advantages in this regard. Cutrale et al.^16^ develop a spectral phasor unmixing algorithm that employed Fourier transform to map spectral images to a two-dimension plane and perform spectral unmixing with clustering methods. Furthermore, Gratton et al.^17^ realize good performance in fluorescent lifetime imaging with spectral phasor analysis. However, these phasor-based nonlinear unmixing methods are still dependent on the performance of clustering methods such as K-Means, though they are analytically easy and computationally fast in phase space.

In recent years, some studies have turned to the use of deep learning for fluorescence microscopy to achieve spectral unmixing. These works use data-driven methods to extract the features of emission spectra instead of utilizating prior emission spectra directly and have advantages of noise insensitivity and good generalization performance. Smith et al.^18^ combine 3-D convolution and Xception^19^ architecture to propose a multi-layer network for the unmixing of fluorescence lifetime imaging. Their network is trained on synthetic datasets and achieves better unmixing results in vivo imaging than that of iterative fitting methods. Li et al.^20^ incorporate the physical process of multi-color fluorescence imaging into the spectral unmixing framework which couples several generator networks to recover the pure images. Manifold et al.^21^ propose an UwU-Net model based on classical U-Net and demonstrate impressive unmixing performance for hyperspectral imaging, mass spectrometry imaging, and Raman scattering imaging. Overall, the deep learning methods have the potential to solve the problems of the prior emission spectra requirement and high computational load in spectral unmixing.

Here, we propose an autoencoder-based fast asymmetric spectral unmixing method (AutoUnmix), which can blindly separate mixed images without reference spectra. Compared with the previous methods that focus on only the spectral unmixing process, our proposed method leverages the physical process of both spectral unmixing and mixing into two networks respectively. Specificially, for the unmixing process in actual microscopy imaging, we design a spectral learning module to learn the mutual information features between channels and a spatial learning module to extract morphological features. In addition, we employ a U-Net^22^ to extract the features of spectral mixing process to further help reconstruct unmixed images accurately. In experiments, we validate that AutoUnmix can separate highly mixed images precisely in around 100ms after training the model on simulated datasets. In addition, a new transfer learning framework is also proposed to further improve the image quality of spectral unmixing by fine-tuning network parameters for the case of real image. On the real-acquired images of mouse cell samples and mouse brain slice samples, we demonstrate that our method has great unmixing reconstruction performance and good generalization. More importantly, AutoUnmix solves the problem that ground truth is not available for the real multi-color fluorescence imaging system and can be applied to different microscopic imaging system without retraining, thus reducing training cost and improving unmixing efficiency.

## 2 Results

### 2.1 Overview of the workflow

We propose a supervised asymmetric autoencoder unmixing network (AutoUnmix) to learn the spectral characteristics of each fluorophore, correct the fluorescent crosstalk from other channels, and output pure images that only contain the corresponding labeled samples for each channel. The AutoUnmix contains the unmixing stage and the mixing stage to imitate the physical process of spectral unmixing and mixing during imaging, as shown in Fig. 1a. In the first stage, the unmixing network learns the features of spectral unmixing by taking mixed images *y* as input and outputting unmixed image 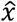. A spectral learning module is employed with channel attention mechanism to learn spectral features between channels and a U-Net is used to extract spatial features. We concatenate and convolve all the generated feature maps in the reconstruction module to obatin the final unmixed output. Then, in the mixing stage, the mixing network extracts the spectral mixing features from the generated unmixed images. Since the physical principle of spectral mixing is not complex, the mixing network adopts a vanilla U-Net to reduce computational complexity and realize fast spectral mixing. Compared to the mixing stage, the unmixing network is more sophisticated. This asymmetric model design helps our method maintain the unmixing performance while reducing the computational cost of the entire model.

**Fig. 1.**
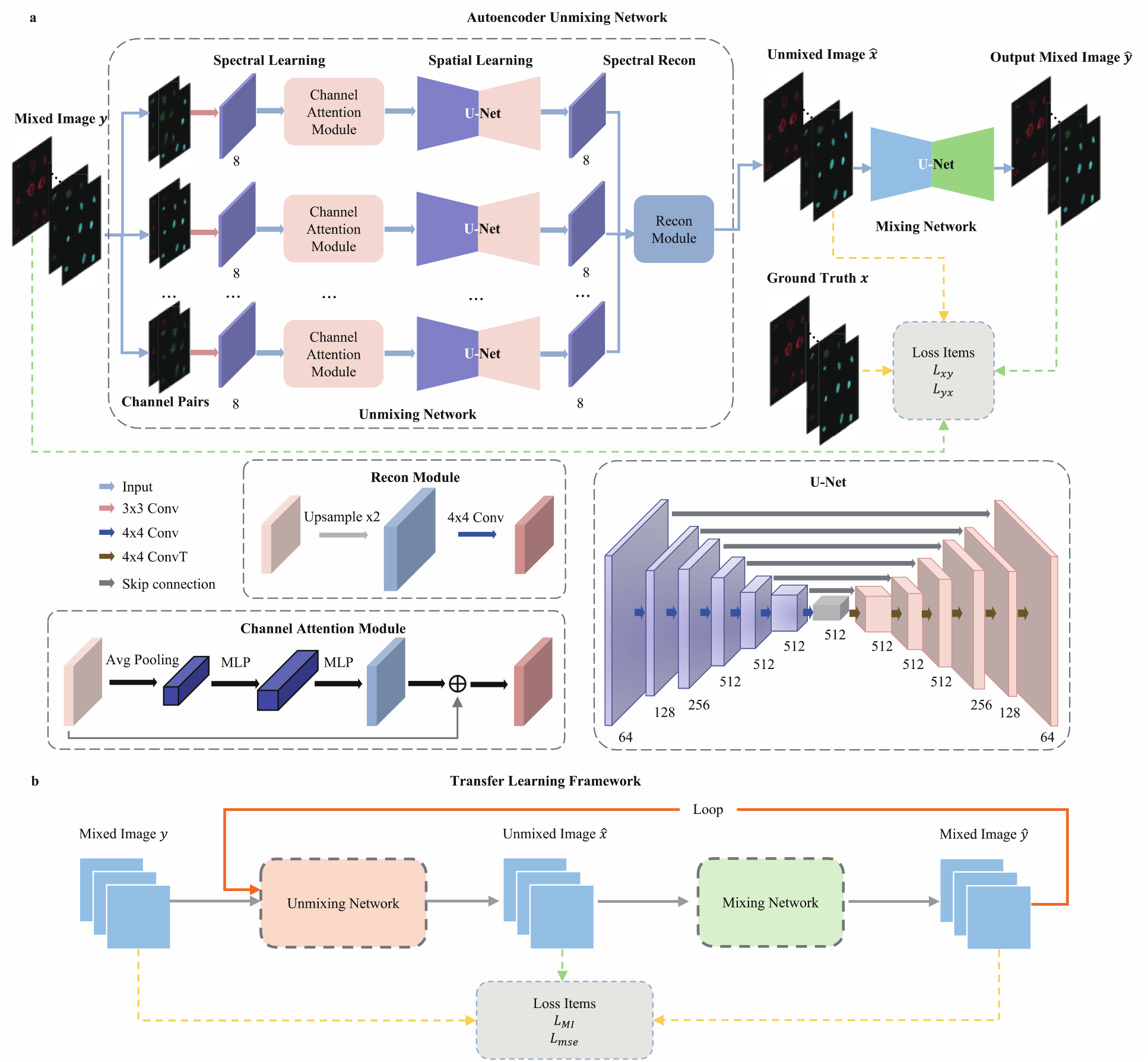
Overview of training procedure and transfer learning procedure of AutoUnmix. **a**. The training procedure of spectral unmixing model. In Unmixing Network, every two channels of input mixed image are selected to extract spectral features by Convolution and Channel Attention Module. A U-Net with shared parameters learns the spatial features in the feature map and then the unmixed image is reconstructed by Recon Module. The backbone of Mixing Network is a U-Net. **b**. The procedure of transfer learning. Mixed image is the initial input. The loop consists of Unmixing Network and Mixing Network. The output from the final loop is the unmixed image for real-acquired mixed images.

In Fig. 1b, a transfer learning framework is designed for the situation that it is impossible to obtain the spectral unmixing ground truth in real experiments. Therefore, the input is the acquired mixed image *y* and the output is the generated unmixed image 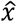. The intermediate result is the mixed image *Ŷ* reconstructed by 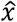. The mixing stage and unmixing stage are iteratively repeated to improve spectral unmixing quality. We design our custom loss function by minimizing MSE and mutual information loss between every two channels. This conforms to the independent and exclusive characteristics of the cell tissues labeled with fluorescence dyes. The design of the loss function also enables us to adjust and optimize the model for different real experimental environments based on the model trained on simulated data, improving the generalization ability and ensuring high-quality spectral unmixing in various situations.

### 2.2 Unmixing performance on simulated datasets

We first compare AutoUnmix with linear unmixing^2^ and LUMoS^14^ in terms of unmixing accuracy on the simulated datasets. To generate simulated data, we select three fluorophores (DAPI, Alexa Fluor 488, MitoTracker Red) with emission spectra highly overlapped and choose three bandpass filters (470/40, 520/40, 600/50nm) corresponding to the spectra peaks of selected fluorophores (Fig. 2a). Then, 345 three-color labeled U-2 OS cell images are artificially synthesized based on the above spectral curves. Each channel of the cell images corresponds to protein, nucleus, and endoplasmic reticulum (ER), respectively. We randomly select 35 images to compare the unmixing performance of linear unmixing, LUMoS, and AutoUnmix (Ours). Linear unmixing is performed according to the theoretical reference spectra (Fig. 2a), while LUMoS does not need emission spectra to unmix images. We quantify the unmixing accuracy by computing the SSIM, PSNR, and MSE between the output unmixed images and the ground truth. Table 1 shows the quantitative evaluation of the unmixing results of the three methods on the synthetic datasets. The performance of linear unmixing and LUMoS methods are closely comparable, with average SSIM values of 0.92 and 0.91 respectively. However, linear unmixing demonstrates significant variance, suggesting a lack of robustness in this approach. Our method achieves an average SSIM of 0.99 on 35 mouse cell images of different shapes, with a small variance, indicating our stable performance and the ability to learn the spectral features of mixed images well. In addition, AutoUnmix completes unmixing within 60ms, which is up to 100 times faster than the other two methods. Fig. 2c shows the unmixing results of the three methods and the ground truth. It is obvious that the unmixing performance of linear unmixing is poor, as it cannot correctly unmix the images in channel 2 and channel 3, and even fails to unmix them completely. In contrast, our method is almost identical to the ground truth. As shown in Fig. 2d, we measured the normalized intensity values along the white arrows in each channel, and our method shows the highest consistency with the ground truth, while linear unmixing and LUMoS do not.

**Table 1.**
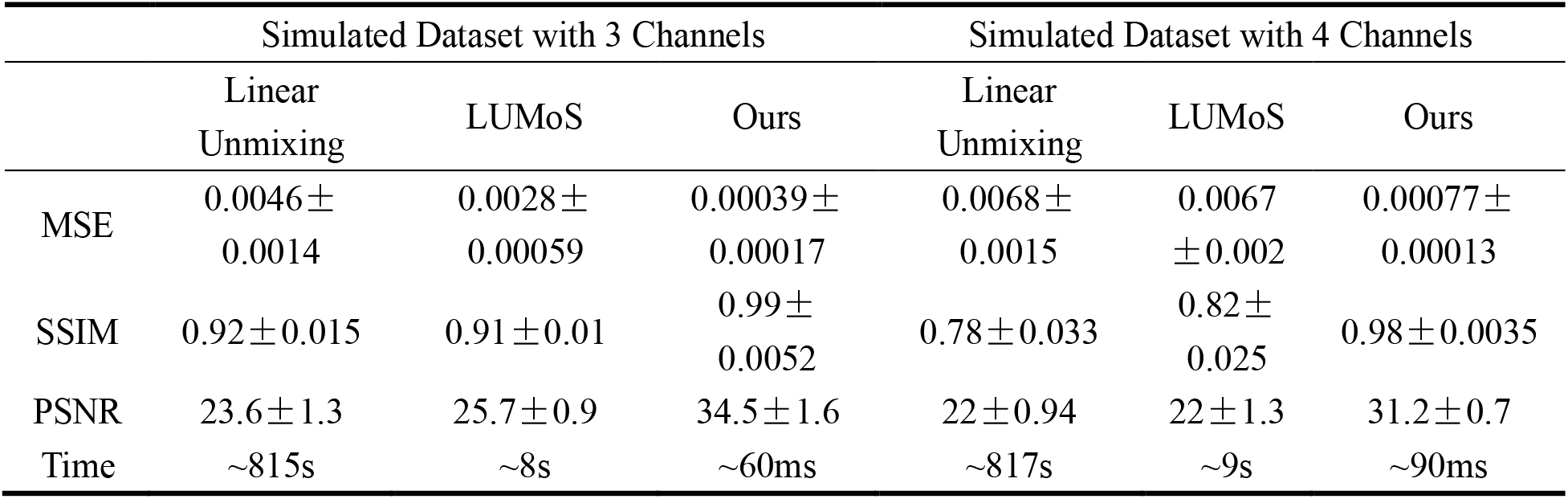
Quantitative comparisons of unmixing performance between the three methods.

**Fig. 2.**
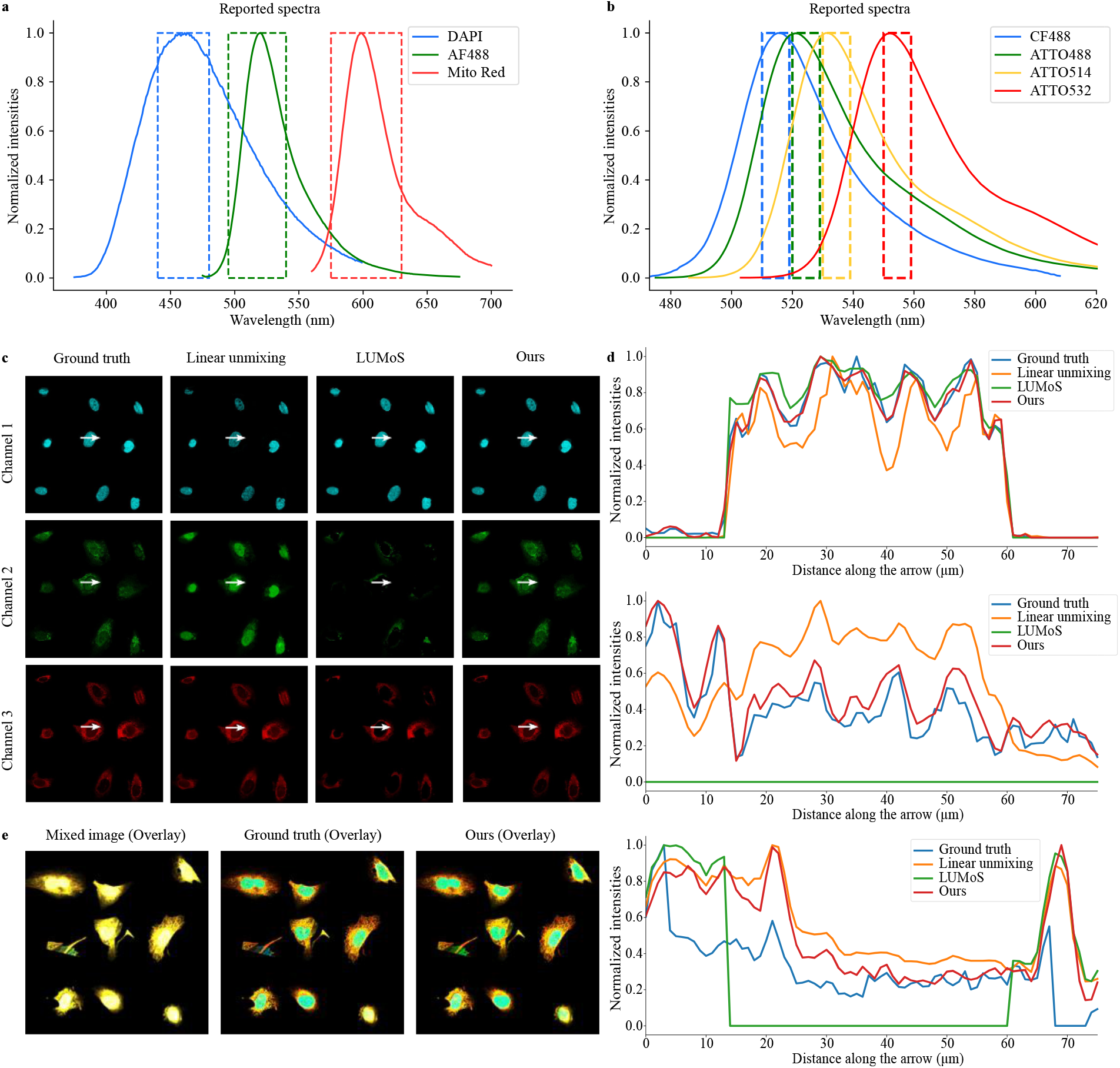
Comparison between linear unmixing, LUMoS and our methods. **a**. The theoretical emission spectral curve of 3 fluorophores (CF488, ATTO514, ATTO532), and the selection of bandpass filter. **b**. The emission spectra of the 4 fluorophores used in the simulated dataset and the detection channels. **c**. The unmixed images of ground truth, linear unmixing, LUMoS and ours. Each row represents channel 1, 2, and 3 respectively. **d**. From top to bottom, normalized intensity values along the white arrows for channel 1, 2 and 3 in **c. e**. Overlay of the results of 4-color unmixing, including the mixed image, ground truth, and the output unmixed image.

Then, we validate the unmixing performance over four dyes highly spectral overlapped: CF488, ATTO488, ATTO514, and ATTO532 (Fig. 2b, bandpass filter of 515/10, 525/10, 535/10, and 555/10nm). A simulated dataset is generated using four-color labeled U-2 OS cell images with each channel corresponding to protein, nucleus, ER, and microtubules. We still use part of the simulated dataset for training and randomly select 35 mixed images for testing. Although the fluorescence dyes have highly overlapping emission spectra, our method (Fig. 2e) still shows good consistency with the ground truth, with an average SSIM of 0.987. In Fig. S1 and S2, we also present the comparison of each channel image between these three unmixing methods as well as the normalized intensity values along the white arrows. The quantified results in Table 1 demonstrate our AutoUnmix can successfully handle various fluorescence mixing situations with fast computational speed.

### 2.3 Validation of blind spectral unmixing

We next validate the ability of AutoUnmix for blind spectral unmixing by choosing fluorophore groups different from those used for training. Our experiments show that our AutoUnmix can successfully separate mixed images with absolutely different emission spectra curves. We select three groups of fluorophores. The emission spectra of the first group (CF488, ATTO514, ATTO532) are similar to the previous four-channel simulated dataset, but it only has three highly overlapped fluorophores. The second group (tagBFP, Cerulean, Citrine) is commonly used for colorful cell imaging, where tagBFP is for nucleus, Cerulean is for cell membrane, and Citrine is for mitochondria. The third group (CF633, Alexa647, CF660R, CF680R) is overlapped relatively balanced. The test datasets are generated based on the three groups of spectral curves (Fig. 3). We directly use the model for 3-channel and 4-channel unmixing trained in section 2.2 to test on the data of the corresponding number of channels. The quantitative results on different combinations of fluorophores are shown in Table S1 and the unmixed images are displayed in Fig. 3. We can see from the overlay outputs that AutoUnmix can separate the mixed images and have good consistency with the ground truth (Fig. S3, S4 and S5 show each channel of the results). Our method achieves SSIM 0.97+ and PSNR 30+ on the generated three-channel mixed images, and SSIM 0.95 and PSNR 28 for the mixed data of four fluorophores. We also evaluate the unmixing results for the same sample image as Fig. 3a generated with largely different ratios of fluorophore combinations in Fig. S3. Whatever the degree of spectral overlapping, our method can still perform well in such tests. The results above show that our AutoUnmix has impressive blind unmixing performance. This indicates our unmixing method enables correctly separating fluorescent dyes with distinct emission spectra from training sets, which is largely attributed to the comprehensive understanding of the underlying physical process of spectral mixing. Consequently, AutoUnmix does not require retraining neural networks for different fluorescent dyes, reducing the heavy cost of re-training while maintaining good unmixing results.

**Fig. 3.**
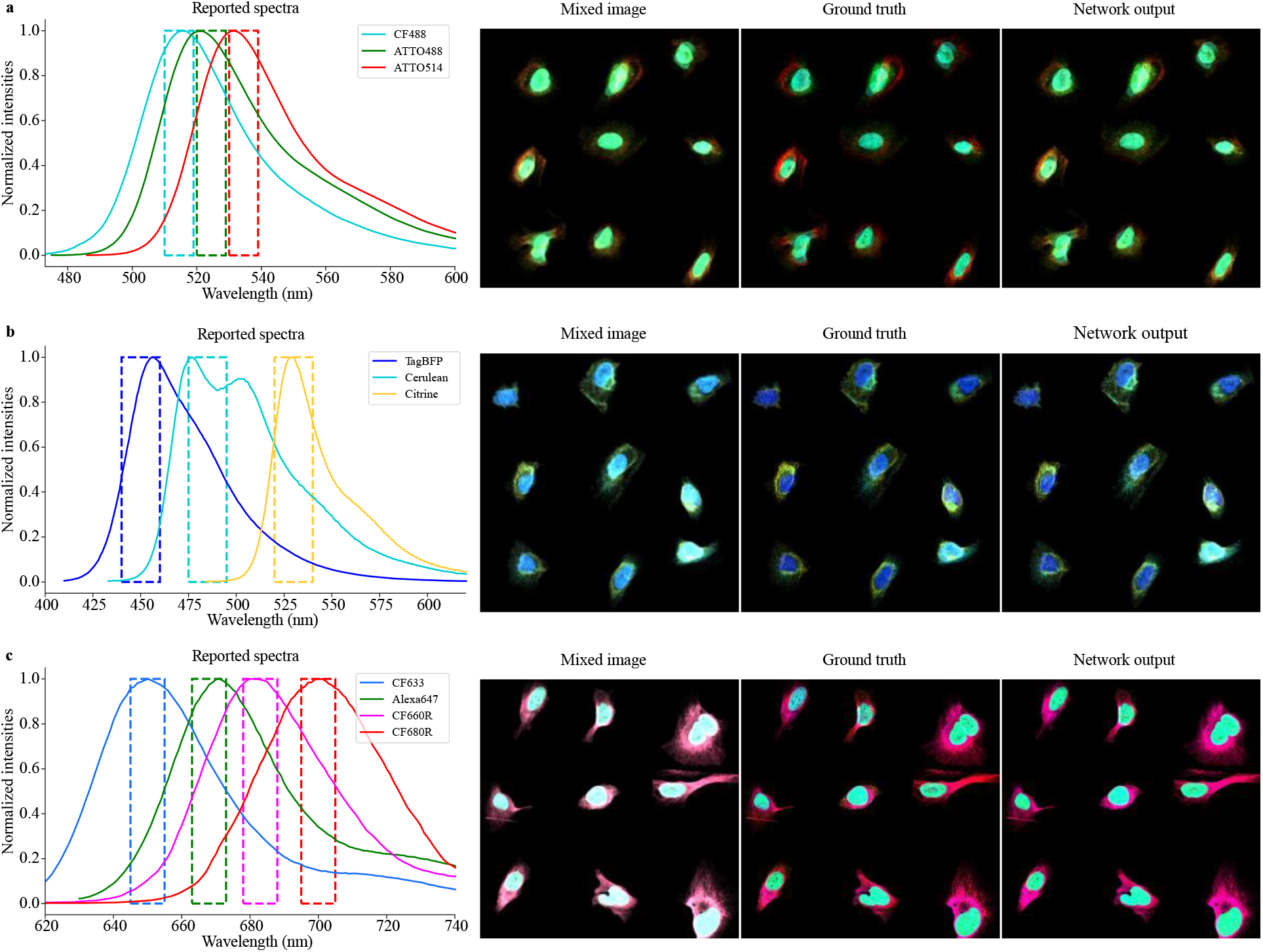
Reference spectra and blind unmixing results of different groups. Different groups of theorectical spectral curve and corresponding bandpass filter. The overlayed images of mixed, ground truth and our output in **a** (CF488, ATTO514, ATTO532), **b** (tagBFP, Cerulean, Citrine) and **c** (CF633, Alexa647, CF660R, CF680R).

### 2.4 Generalization performance on real biological samples

In real biological experiments, it is impossible to obtain the ground truth of spectral unmixing, which imposes a great challenge to supervised deep learning methods. Here, we propose a transfer learning framework (Fig. 1b) to remedy the absence of ground truth by applying finetuning. First, we train a base model using the simulated datasets generated from reference spectra of the same number of fluorophores. Then we initialize the parameters of AutoUnmix with the base model and iteratively finetune the parameters on the real-acquired images to learn the distribution gap between synthetic data and real data. The finetuning is stopped when the MSE loss between input mixed image and output mixed image is smaller than a threshold. We demonstrate the unmixing performance of our proposed method on three real biological samples.

We tested the mouse cell samples with DAPI, F-Actin and MitoTracker Red. The 3-channel mixed images were obtained using a TIRF microscope. More details are in Methods. We employ the base model in section 2.2 where the spectral curve for training is quite similar to that of mouse cell samples and then finetune the model on the real acquired mixed images. Fig. 4a displays the original mixed image, linear unmixing results, LUMoS results, and Ours. In channel 1, the nucleus structure cannot be completely reconstructed by linear unmixing method, and LUMoS can maintain the cell structure to some extent while losing some clear details. In channel 2, crosstalk from channel 1 is visible in the mixed image. Only our method can remove all nuclei while reducing the noise signal in the image simultaneously. In channel 3, it can be seen that LUMoS is indeed able to remove the crosstalk from channel 1 and 2, but destroys the mitochondrial structure. However, our method can both maintain the mitochondrial structure and remove the crosstalk from the nuclei, indicating that AutoUnmix can indeed separate real mixed images.

**Fig. 4.**
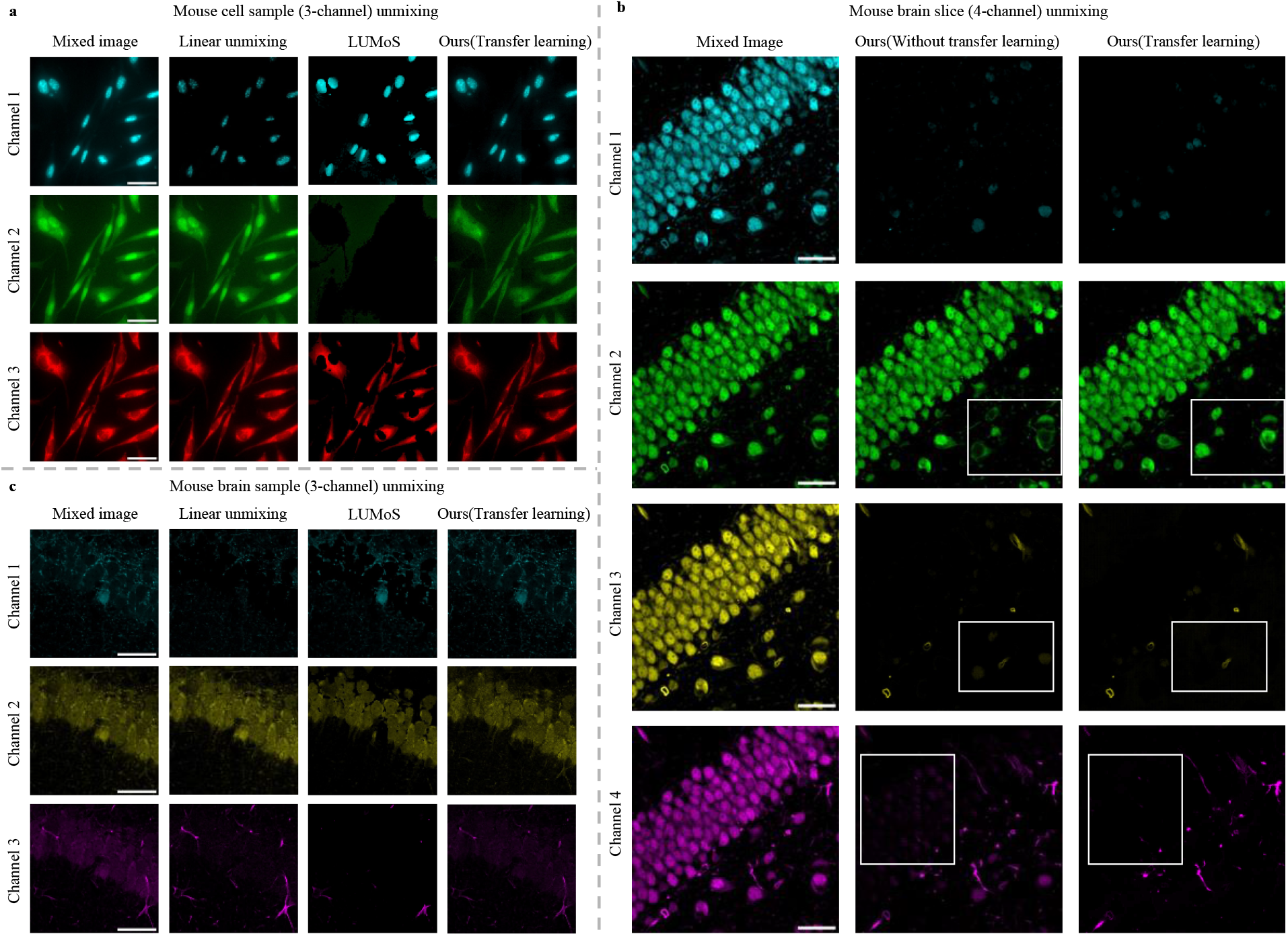
Unmixing results on real biological samples. **a**. Unmixing results on 3-channel mouse cell samples exhibiting similar spectra and morphology to training data. Each column is results of the three methods and each row represents different spectral channel. Scale bar: 50μm **b**. Unmixing results of the AutoUnmix with and without transfer learning on 4-channel mouse brain slice samples with different morphology. Scale bar: 30μm **c**. Unmixing results on 3-channel mouse brain slice samples with different morphology and emission spectra. Scale bar: 40μm.

Next, we verify the unmixing quality of AutoUnmix on samples exhibiting different morphology. Fig. 4b displays the mixed images of mouse brain slices with highly overlapped spectra (CF488, ATTO488, ATTO514, ATTO532), which shows quite different morphological structure from training datasets. Compared with linear unmixing and LUMoS, AutoUnmix with transfer learning can achieve better unmixing performance as shown in Fig. S6. Our method without transfer learning cannot completely remove some of the cell structures that crosstalk into channel 1. As a contrast, by employing transfer learning framework, more cell details can be recovered in the white rectangle in Fig. 4b, indicating that our transfer learning framework can significantly improve the output quality.

We further test the proposed method for samples with different morphology and emission spectra as shown in Fig. 4c. The three fluorophores are CF488, ATTO514, and ATTO532. We directly use the previously trained network to finetune on the real-acquired images, and the impressive unmixing results can be obtained. Although the three methods can remove the crosstalk from channel 2 into other channels, linear unmixing loses a lot of tissue structure in channel 1, and LUMoS lacks some details in channel 3, while our method maintains the structure of the cellular tissue well. These results show that our method can still maintain high unmixing performance for samples with different morphologies by performing transfer learning on real sample images. This demonstrates AutoUnmix does not need to retrain networks for real experimental conditions or rely on reference spectra.

## 3 Discussion

In this work, we propose a new deep learning method for fluorescence spectral unmixing, termed AutoUnmix, as an alternative flexible choice for separating mixed images. Our proposed method designs an asymmetric autoencoder model that incorporates the process of spectral mixing and unmixing as two stages in deep neural networks to simultaneously learn spectral characteristics during this procedure. On simulated datasets with 3- and 4-color mixed images, we can achieve SSIM of 0.99 and PSNR of 30+, which show great superiority over traditional linear unmixing methods and machine learning-based LUMoS method.

Further, we demonstrate the unmixing performance of our AutoUnmix with the proposed transfer learning framework. By fine-tuning the network on single real-acquired image, our method successfully unmixes images on both real mouse cell samples and mouse brain slice samples that have inconsistent morphology with training sets and largely different emission spectra of labeling fluorophores from the theoretical spectra. Compared to the other two unmixing methods, our method shows impressive unmixing performance and generalization ability. This result highlights that we present a novel approach to address the issue that the ground truth of real samples is not available and training images differ from real biological images. This also can be attributed to the autoencoder architecture in AutoUnmix that is naturally applicable to network training without ground truth and improves unmixing performance by iterative looping between the encoder and decoder. To some extent, our approach solves the problem that the independent identically distribution assumption between synthetic data and real data is not satisfied. However, it is noted that since our method dose not learn morphological features during training, some channels of real samples cannot be absolutely separated, which deserves further improvement in subsequent research.

Finally, we aim to extend the application of our AutoUnmix to various unmixing imaging modalities. Our transfer learning method can further adapt to different real imaging systems and bio-samples labeled with different dyes, such as mRNA imaging^23–25^, cell tracing^26^, fluorescent barcoding techniques^13,27^, multidimensional cytometry^28^ and sub-cellular structures locations^29^. Moreover, our unmixing network architecture can be potentially used in label-free detection of intracellular structures^21,30^. Our method may also perform well in biomedical tasks such as virtual staining^31–33^, as they also involve multi-spectral imaging task. Additionally, our method can be combined with spectral phasor methods^17,34,35^ to achieve spectral unmixing while utilizing the computational efficiency in the Fourier space. We can also explore replacing modules in our mixing network and unmixing network with more efficient modules like vision transformer blocks^36, 37^ to better extract spatial features and address deficiencies on unseen morphologies. Overall, we anticipate that our AutoUnmix will be useful in medical imaging, microscopy imaging, remote sensing^38,39^, and an even broader range of computer vision tasks.

## 4 Materials and methods

### Data Acquisition

#### Simulated datasets

Although some existing methods, such as PICASSO using antibody complex formation techniques, can acquire images with almost no spectral crosstalk, such methods involve tedious biological sample preparation process. Here, we generate datasets with various fluorophores and spectra profiles to simulate different imaging environments, which are then used for training and testing. After that, we apply the trained model to unmixing tasks for real sample images. The generation procedure is shown in Fig. S7. Firstly, we select 345 4-channel images of U-2 OS cells from the DULoc^40^ dataset as ground truth, which were obtained from the HPA^41^ database and processed by the cell masks tool. Each image has 9 cells and 4 channels, where each channel corresponds to protein, nucleus, ER, and microtubule. Second, we select multiple fluorophore groups and download the reference spectra curve of them from FPBase^42^ database, including DAPI, AF488, CF488, AF532, and so on. We also set the center wavelength and witdth of bandpass filters according to the selected fluorophores. The intensity values of each channel for each pixel of the ground truth image are mixed using linear mixing, and then additional Poisson noise is added to each channel to simulate the shot noise.

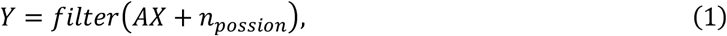

where *Y* represents generated mixed image, *X* is the original unmixed image, *A* is the mixing matrix, *n*_*poisson*_ is Poisson noise, and *filter* represents Gaussian filter and median filter. Images are convolved with a Gaussian filter with a standard deviation of 0.5 and a 3×3 median filter to represent the real-world diffusion effect to obtain mixed images. The purpose of this workflow is to closely simulate the real conditions that biological samples often exhibit overlapping tissue structures. The spatial dimensions of each synthesized image are 512×512, and 3- and 4-channel images are generated, respectively.

#### Real biological sample acquisition

Real biological experiments were performed on mouse cells stained with DAPI for nuclei, F-Action for actin fibers, and MitoTracker Red for mitochondria. Multicolor images (Fig. S7a) were obtained using a TIRF microscope, Nikon ECLIPSE Ti2-E. The fluorophores were excited with three lasers at 405, 488, and 561nm, respectively. The size of the acquired mixed images is 512×512×3. Then, we normalize each channel of the images to facilitate the subsequent unmixing tasks. We also used 3-color (CF488, ATTO514, ATTO531) and 4-color (CF488, ATTO488, ATTO514, ATTO532, Fig. S7a) fluorescence images of mouse brain slices^10^ to test the unmixing performance of our method on real samples that are not consistent with the distribution of the training datasets.

### Network details

The unmixing network, as shown in the left part of Fig. 1a, takes the mixed image *y* with dimension (H, W, C) as input, where H and W denote the spatial size of *y*, and C denotes the number of channels of *y*. We design a Spectral Learning Module (SLM) containing a convolution and a Channel Attention Module (CAM)^43^. The operation of CAM can be formulated as:

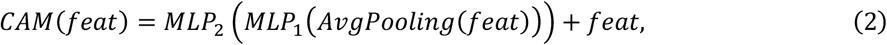

where *feat* denotes the high-dimension feature map, *MLP*_1_ containing a fully connected layer maps the input of size 1 × 1 × *C*_*feat*_ to size of 1 × 1 × *C*_*feat*_ / 16 and *MLP*_2_ is also a fully connected layer performing the inverse operation of *MLP*_1_. The SLM sequentially selects each channel pair *i* and *j* and generates the corresponding high-dimension feature maps as follows:

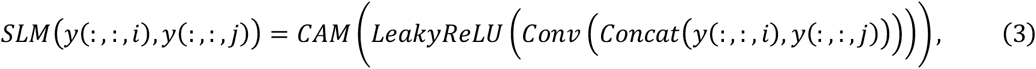

where *y*(:, :, *i*) and *y*(:, :, *j*) denote the corresponding channel *i* and *j* of mixed image *y*. Each feature map extracted by SLM is then fed into a U-Net that shares parameters to reduce computational cost to learn the spatial features. Finally, in the spectral reconstruction module, all of the U-Net outputs are concatenated and convolved to obtain the unmixed image 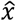, which can be expressed as:

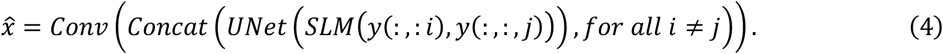

The mixing network, as shown in the right part of Fig. 1a, takes the unmixed image 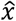 generated in the previous stage as input and employs a lightweight U-Net to recover the mixed image.

To ensure better reconstruction image quality, we reduce the difference between the output unmixed image and ground truth while preserving the morphological structure of tissue cells. The loss function during training includes Mean Squared Error (MSE) loss and Structural Similarity Index Measure (SSIM)^44^ loss, shown as follows,

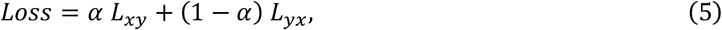

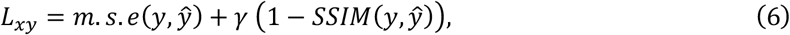

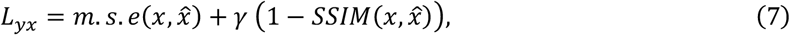

where m.s.e represents the MSE loss, *L*_*xy*_ is the loss term guiding the mixing network training and *L*_*yx*_ is the loss term guiding the unmixing network training. *α* controls the relative importance between *L*_*xy*_ and *L*_*yx*_ and it can be selected empirically. Here, we set it as 0.5.

The loss function in transfer learning is different from the one in the training with simulated data. In addition to computing the MSE loss between the input and output mixed images, mutual information loss^45^ also needs to be computed.

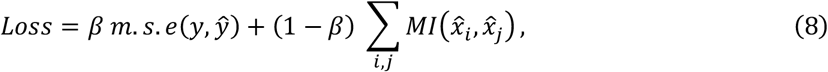

where *β* represents the coefficient, and the latter term represents the loss calculating the mutual information between every two channels and summing them up. Specificially, the mutual information between two gray scale images is formulated as follows:

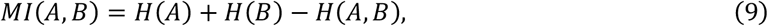

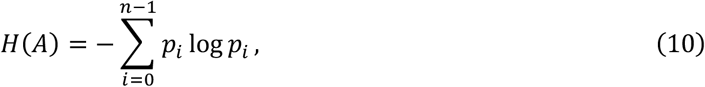

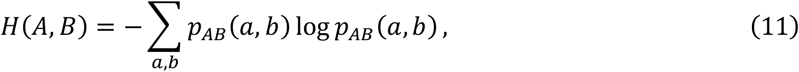

where *n* denotes the number of gray values in the image, usually taken as 256 for 8-bit grayscale images, *p*_*i*_ denotes the probability that a pixel point with a gray value *i* is present in that image, and *p*_*AB*_(*a, b*) is the joint probability with gray-scale value *a* in channel A and gray-scale value *b* in channel B.

### Metrics

Following the numerical evaluation methods proposed in the previous papers, the SSIM, PSNR and MSE are used to evaluate the results of the spectral unmixing methods. The values of the SSIM metrics are between 0 and 1. The higher SSIM and lower MSE values represent a high-quality output result. MSE is a general objective evaluation index in measuring image quality, while SSIM assures a high degree of structural consistency between output and ground truth. The SSIM is defined as

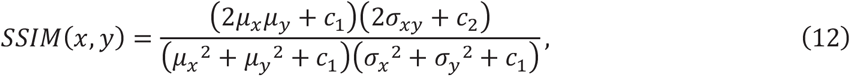

where *μ*_*x*_, *μ*_*y*_ and *σ*_*x*_^2^, *σ*_*y*_^2^ are the averages and variances of images *x, y*; *σ*_*xy*_ is the covariance of *x* and *y*; *c*_1_, *c*_2_ are the variables used to stabilize the division with a small denominator. We also use PSNR to measure the ratio between the maximum possible power of a signal and the power of corrupting noise that affects the fidelity of its representation. The higher PSNR represents better image reconstruction quality. PNSR is defined as follows based on MSE:

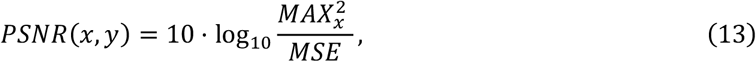

where MSE is the mean square error between image *x* and image *y, MAX*_*x*_ is the maximum possible gray-scale value of the image, which is 255 in our experiments.

### Network training and testing

The model used in this paper is trained on the simulated datasets using randomly initialized parameters and optimized using our custom loss function with stochastic gradient descent. Our proposed spectral unmixing framework utilizes the Adam optimizer with the learning rate of 0.0002, the beta of 0.5 and 0.999, and is trained for 200 epochs. The simulated datasets are split into training and testing sets in a 4:1 ratio. All the images are cropped into a patch size of 256×256 and augmented using rotations and flips to enlarge training sets. All the training and prediction are performed on a server running Ubuntu 18.04, equipped with an Intel Xeon 4216 processor and Nvidia RTX 2080Ti GPU. On our machine, training the 3-channel and 4-channel unmixing model takes approximately 3 and 5 hours respectively, while the prediction time for a single test image sized of 256×256 is around 100 milliseconds.

## Code availability

Our code of AutoUnmix is available from the corresponding author upon reasonable request.

## Data availability

The data that support the findings of this study are available from the corresponding author upon reasonable request.

## 5 Acknowledgements

This work was supported in part by the National Natural Science Foundation of China (62031023), in part by the Shenzhen Science and Technology Project (JCYJ20200109142808034 & GXWD20220818170353009 & JSGG20191129110812708), and in part by Guangdong Special Support (2019TX05X187).

