## Supplementary Information for "AutoUnmix: an autoencoder-based spectral unmixing method for multi-color fluorescence microscopy imaging"

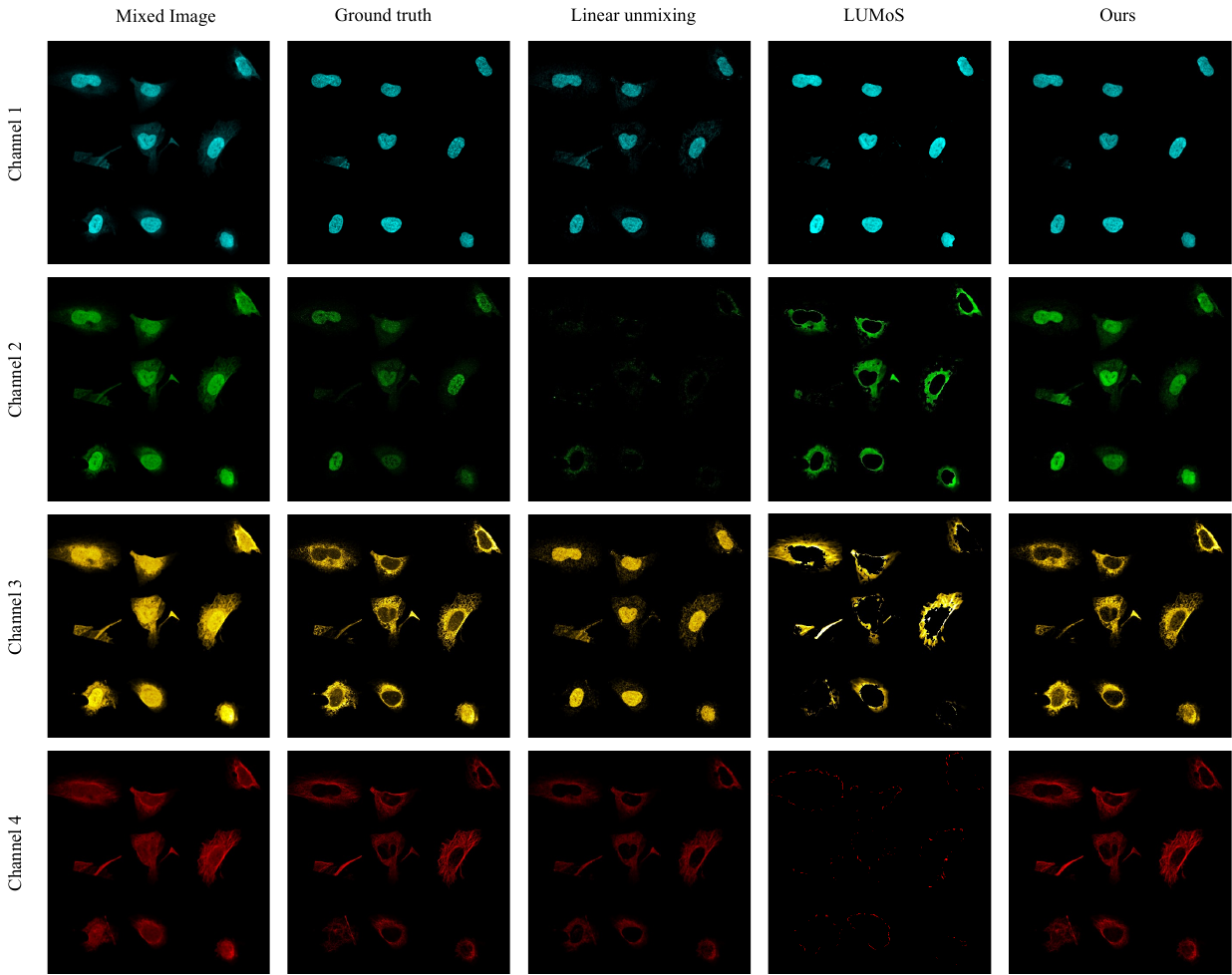

**Fig S1 Comparison between linear unmixing, LUMoS, and Ours on 4-channel simulated data of Fig. 2e** Each row corresponds to each channel of the images. Columns are mixed image, ground truth, results of linear unmixing, results of LUMoS, and results of Ours. Linear unmixing: MSE 0.00578, SSIM 0.786, PSNR 22.4. LUMoS: MSE 0.00769, SSIM 0.77, PSNR 21.2. Ours: MSE 0.000749, SSIM 0.985, PSNR 31.3

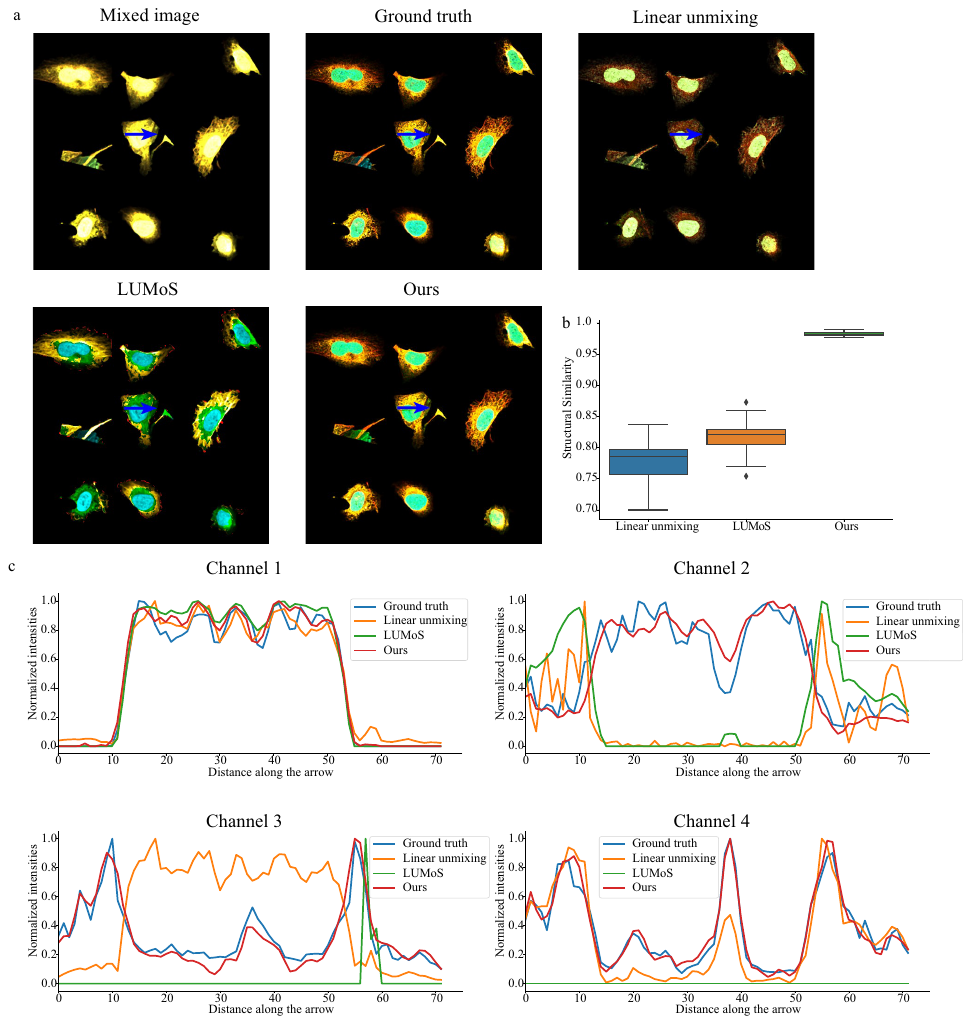

**Fig S2 Quantitative comparison between linear unmixing, LUMoS, and Ours on the 4-channel simulated data** **a.** Overlay of the mixed image, ground truth, linear unmixing, LUMoS, and Ours in Fig. S1. **b.** Boxplot of SSIM results of linear unmixing, LUMoS, and Ours on the 4-channel simulated data. Our method has highest SSIM with smallest variance. **c.** Normalized intensity values along the blue arrows for each channel in **a**, which shows that our method has high consistency with ground truth compared to other two methods.

**Table S1 Blind unmixing performance on datasets with different combinations of fluorophores**

|  | MSE | SSIM | PSNR |
| --- | --- | --- | --- |
| Group 1 (Fig.3a) | 0.0092±0.0013 | 0.97±0.0063 | 30±1.3 |
| Group 2 (Fig.3b) | 0.0085±0.0012 | 0.97±0.006 | 31±1.2 |
| Group 3 (Fig.3c) | 0.011±0.0014 | 0.95±0.013 | 28±0.95 |

The quantitative results on different combinations of fluorophores are shown in Supplementary Table S1corresponding to Fig. 3. Our method achieves SSIM 0.97+ and PSNR 30+ on the generated 3-channel mixed images, and SSIM 0.95 and PSNR 28 for the mixed data of four fluorophores,. The results indicate AutoUnmix has great potential for impressive generalization performance.

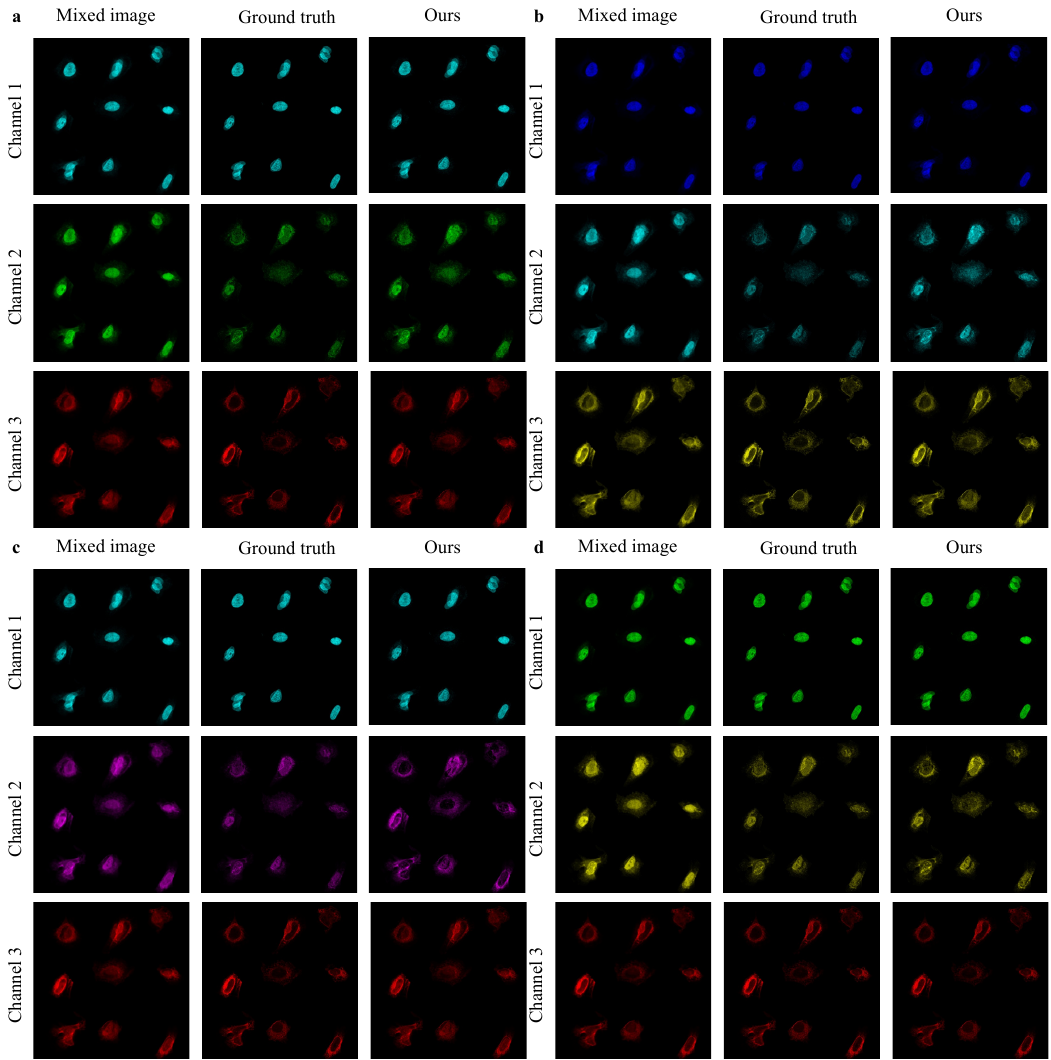

**Fig. S3 Results of blind unmixing for Fig. 3a.** Each channel of the results is shown in the figures above. We show the unmixing results for the same sample image as in Fig. 3a generated with largely different relative ratio of fluorophore combinations (**a**: CF488, ATTO488, ATTO514, bandpass filter: 515/10, 525/10, 535/10nm; **b**: TagBFP, Cerulean, Citrine, bandpass filter: 450/20, 485/20, 530/20nm; **c**: CF633, CF660R, CF680R, bandpass filter: 650/10, 683/10, 700/10nm; **d**: AzamiGreen, Citrine, mCherry, bandpass filter: 505/20, 530/20, 600/20nm;). All of these images are successfully unmixed with our method with SSIM 0.99+ and PSNR 34+, even under significant different relative ratios.

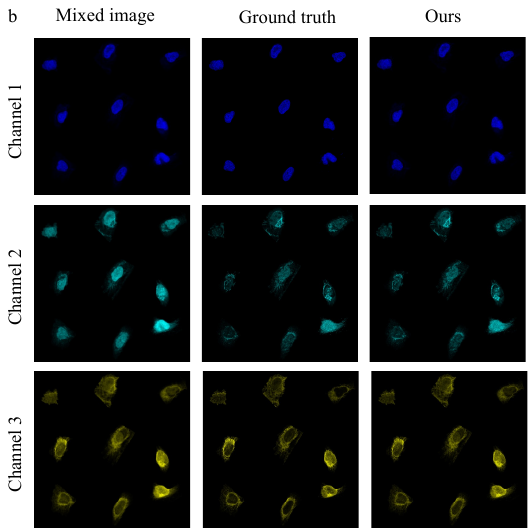

**Fig. S4 Results of blind unmixing for Fig. 3b.** Each channel of the results is shown in the figures above. The mixed image shows channel 2 and channel 3 are highly overlapped and is consistent with the spectral curve in Fig. 3b. Our unmixing output can address such spectra leaks, leading to high consistency with ground truth.

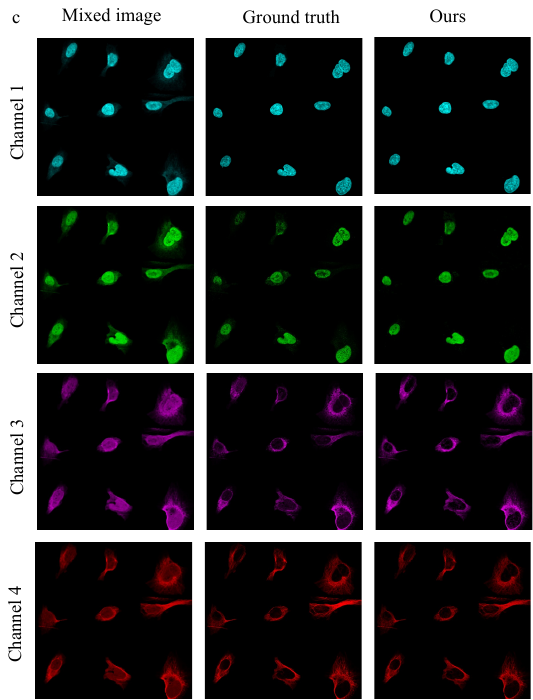

**Fig. S5 Results of blind unmixing for Fig. 3c.** Each channel of the results is shown in the figures above. The mixed image shows channel 2, 3 and 4 are highly overlapped. Compared with ground truth, our results are almost the same and separate different cell structures explicitly.

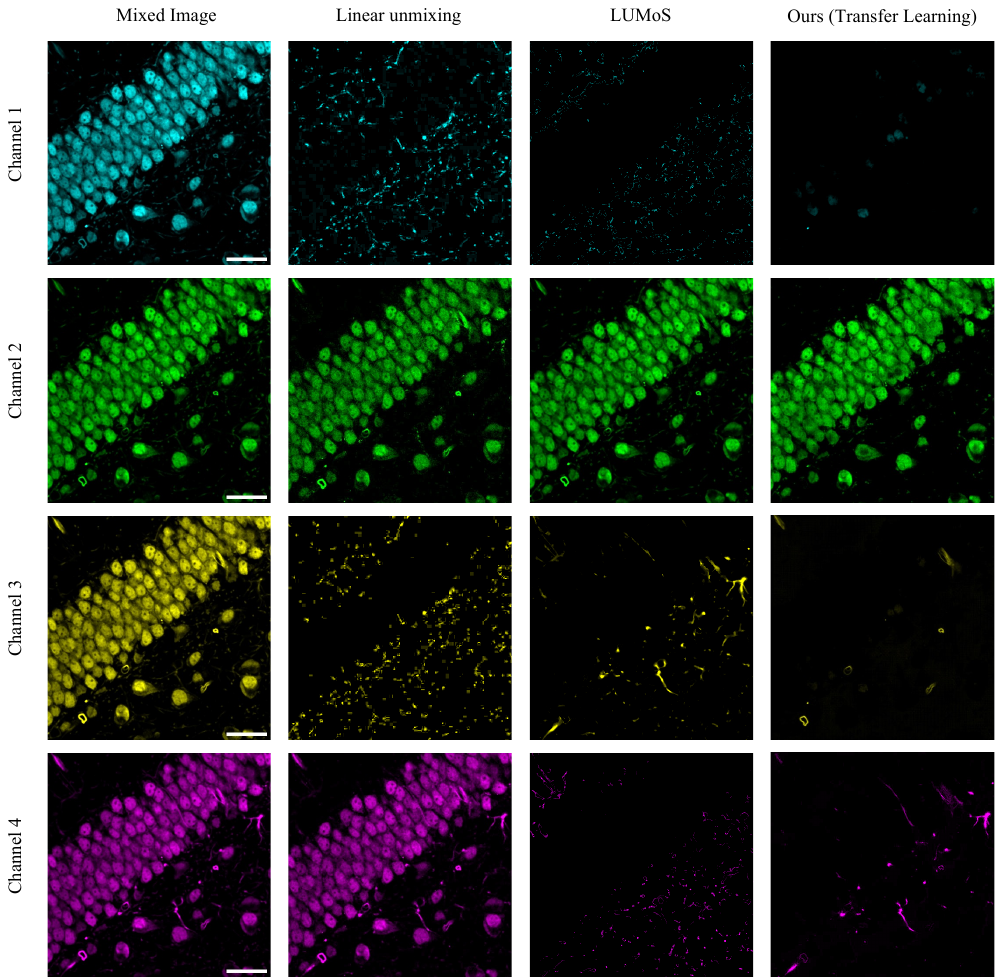

**Fig. S6 Comparison between linear unmixing, LUMoS, and Ours on the real 4-channel sample in Fig. 4b.** Each row shows the unmixing results of each channel between three methods. Scale bar: 30μm. Our method with transfer learning can achieve best unmixing performance. Linear unmixing cannot separate the sample from original highly overlapped images, while LUMoS can separate them but cannot reconstruct complete sample structure in channel 1 and 4.

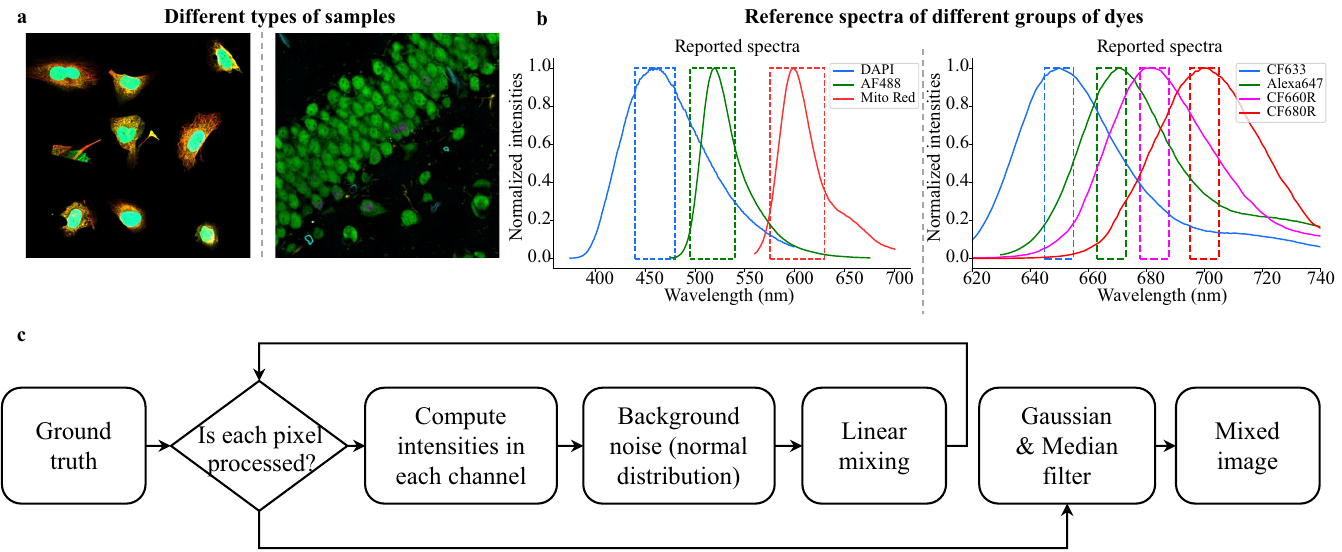

**Fig. S7 Generation flow of simulated datasets. a.** Different multi-channel images selected as ground truth. **b.** Reference spectra of different combinations of dyes. **c.** The generation flow of simulated datasets that translates ground truth into mixed images with the selected sample images and reference spectra using linear mixing. The generation flow corresponds to the description of Simulated datasets of Data Acquisition in Methods.

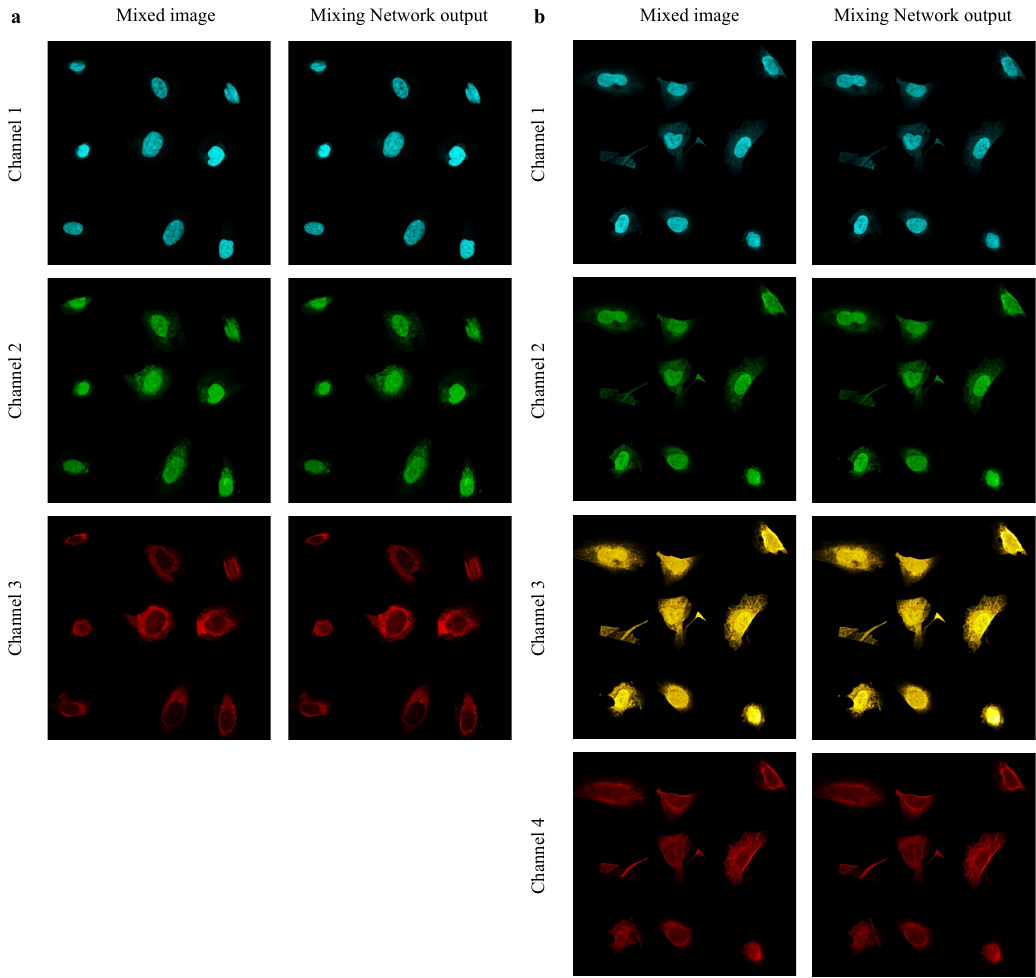

**Fig. S8 Results of mixing network. a.** 3-channel original mixed image and the output mixed image from the mixing network of our AutoUnmix. MSE: 0.000111, SSIM: 0.998, PSNR: 39.6. **b.** 4-channel original mixed image and the output mixed image from our AutoUnmix. MSE: 0.000171, SSIM: 0.997, PSNR: 37.7. The figure and the quantitative results demonstrate that our method can learn from the spectral mixing process and reconstruct high quality mixed image.

**Table S2 Unmixing performance on simulated datasets between U-Net and Ours**

|  |  | SSIM | PSNR |
| --- | --- | --- | --- |
| Group 1 | U-Net | 0.993±0.0023 | 34.3±1.15 |
|  | Ours | **0.997±0.0052** | **34.5±1.56** |
| Group 2 | U-Net | **0.98±0.0029** | 31.1±0.77 |
|  | Ours | 0.98±0.00035 | **31.2±0.7** |
| Group 3 | U-Net | 0.97±0.0065 | 29±1.3 |
|  | Ours | **0.97±0.0063** | **30±1.3** |
| Group 4 | U-Net | 0.97±0.007 | 30±1.3 |
|  | Ours | **0.97±0.006** | **31±1.2** |
| Group 5 | U-Net | 0.94±0.012 | 26±0.77 |
|  | Ours | **0.95±0.013** | **28±0.95** |

To validate that our proposed unmixing network can indeed improve unmixing performance, we also replace the unmixing network in our method with a U-Net. The quantitative results are shown for different simulated datasets tested in our work. Group 1 and group 2 correspond to the 3-channel unmixing and 4-channel unmixing in Fig. 2 respectively. Group 3, 4 and 5 correspond to the blind unmixing validation for different reference spectra in Fig. 3, respectively. We can clearly conclude that our method outperforms the U-Net method in all simulated datasets, though U-Net can achieve close unmixing results to Ours in Group 1 and 2.

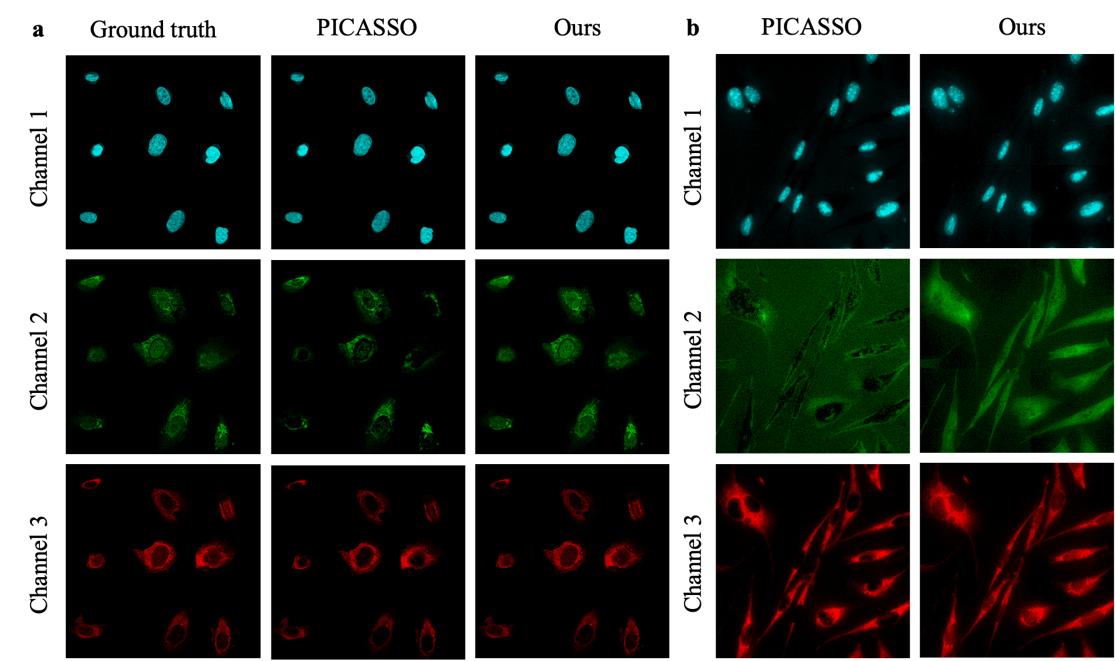

**Fig. S9 The visual comparison between ours and PICASSSO. a.** The unmixed images of ground truth, PICASSO and ours on 3-channel simulated datasets. Each row represents channel 1, 2, and 3 respectively. Both ours and PICASSO can reconstruct images in channel 1. However, PICASSO cannot maintain the cell structure in channel 2 and lose some cell edges in channel 3. Our method performs high consistency with the ground truth. **b.** The unmixed results for real-acquired images comparing PICASSO and ours. The PICASSO method fails to reconstruct the complete structure in channel 2.

**Table S3 Quantitative comparison between PICASSO and Ours**

|  | MSE | SSIM | PSNR | Time |
| --- | --- | --- | --- | --- |
| Ours (AutoUnmix) | 0.0003 | 0.999 | 35.4 | ~60ms |
| PICASSO | 0.0009 | 0.969 | 30.5 | 6.5s |

The quantitative comparisons are listed in the table above, which indicating that our AutoUnmix surpasses PICASSO with a large margin. In addition, our AutoUnmix can achieve unmixing within 60ms with the help of RTX 2080 GPU, while PICASSO requires 6.5s to finish this task, which corresponds to the 100-fold improvement in our manuscript. The results in this table and Fig. S9 demonstrate that our AutoUnmix has superior performance over PICASSO in terms of unmixng image quality and algorithm efficiency.
